## Supplementary Figures for "High-performance chemical and light-inducible recombinases in mammalian cells and mice"

**SUPPLEMENTARY FIGURE LEGENDS**

**Supplementary Figure 1.** **HHPred alignment of VCre to Cre.**

Results from VCre HHPred alignment to Cre recombinase (PDB: 1XO0, chain A) are reproduced here with additional annotations indicating split locations, catalytic residues, and secondary structure names adapted from a literature source^35^. Three-dimensional structure of Cre recombinase is plotted with beta-sheets annotated in blue and alpha helices in pink.

**Supplementary Figure 2.** **Cloning strategy for building split recombinases.**

(**a**) A schematic showing the process for cloning split recombinase fragments into CID backbones. (**b**) Series of vectors containing CID and LID domains. NLS indicates the SV40 nuclear localization sequence.

**Supplementary Figure 3.** **Requirements of four gibberellin-inducible split Flp recombinase systems.**

(**a**) Flp was split into two fragments at amino acid S and fused to gibberellin-inducible (GAI/GID1) dimerization domains. Split recombinases were transfected along with reporters that yield green fluorescence protein (GFP) expression upon site-specific recombination (**b**) Constitutive Flp (black square) or a blank vector (gray square) were transfected to indicate highest or lowest expected GFP expression. Split Flp recombinase components (N-terminal or C-terminal halves) fused to chemical inducible dimerization (CID) domains were transfected in all combinations and drug (colored shapes) or no drug (white shapes) was added to the cell culture medium after two hours. (**c**) Flp was split into two fragments at amino acid S with no fusion to dimerization domains. Split Flp recombinase components were transfected along with reporters that yield green fluorescence protein (GFP) expression upon site-specific recombination. (**d**) Constitutive Flp (black square) or a blank vector (gray square) were transfected to indicate highest or lowest expected GFP expression. Split Flp recombinase components (N-terminal or C-terminal halves) domains were transfected in all combinations. M.F.I. indicates mean fluorescence intensity measured with arbitrary units (a.u.). Errors bars represent arithmetic standard error of the mean of three transfected cell cultures.

**Supplementary Figure 4.** **Comparison of original split Cre (diCre) with newly developed split Cre recombinases.**

(**a** and **b**) Cre was split into two fragments at amino acid S and fused to gibberellin-inducible (GAI/GID1) heterodimerization domains at various termini. Split recombinases were transfected along with reporters that yield green fluorescence protein (GFP) expression upon site-specific recombination (**c**). Constitutive Cre (black square) or a blank vector (gray square) were transfected to indicate highest or lowest expected GFP expression. Split recombinases were also transfected and drug (blue squares) or no drug (white squares) was added to the cell culture medium after two hours. M.F.I. indicates mean fluorescence intensity measured with molecules of equivalent fluorescein (MEFL). Error bars of M.F.I. indicate the geometric standard deviation of means between three transfected cell cultures.

**Supplementary Figure 5. Benchmarking Cre, Flp and ɸC31 systems performance over time using mean fluorescence intensity (M.F.I.), fold activation (Fold), and signal-to-noise ratio (SNR).**

(**a**) A schematic showing the incorporation of three chemical inducible dimerization (CID) systems or nuclear shuttling mutated estrogen receptor (ER^T2^) system. (**b**) An SNR metric can be used to capture distinguishability of “on” and “off” states, accounting for both the absolute difference in mean signal expression and spread (“noise”) of the distributions. (**c**) White or colored squares represent M.F.I. for minus drug (or blank vector) and plus drug (or constitutive recombinase), respectively. Fold activation and SNR are represented by circles and diamonds, respectively. The three metrics, M.F.I. (**d**), Fold (**e**), and SNR (**f**), were captured over 100 hours post-transfection for particular amino acid splits of inducible Cre, Flp and ɸC31 systems incorporating the gibberellin (GIB), rapalog (RAP) and abscisic acid (ABA) dimerization domains GAI/GID1, FKBP/FRB and PYL/ABI, respectively. A 4-hydroxytamoxifen (4OHT)-inducible pCAG-ER^T2^-Cre-ER^T2^ construct is also included in (**b**).

**Supplementary Figure 6. Single-cell fluorescence representation of time-course data.**

Single-cell data represented as smoothed scatterplots are plotted for the 21 hour (**a**), 52 hour (**b**) and 100 hour (**c**) time points captured over the 100 hour time-course of the experiment in **Figure 2b**. Axes represent the expression of transfection marker, BFP, in molecules of equivalent fluorescein (MEFL) and GFP output, in MEFL, from recombinase reporters. Density of cytometric events are represented ranging from no events (dark blue) to highest density in dark red. White dots represent single event outliers. Events under a BFP MEFL cutoff of 1x10^6^ (representing non-transfected or poorly transfected cells) were removed from analysis and a red line at GFP MEFL = 1x10^6^ indicates a boundary between cells that are “OFF” (below 1x10^6^) and cells that are “ON” (above 1x10^6^).

**Supplementary Figure 7. Benchmarking Cre systems performance over time using binned mean fluorescence intensity (M.F.I.), fold activation (Fold), and signal-to-noise ratio (SNR).**

(**a**) Legend indicating labels for time-course values. GFP M.F.I., Fold, and SNR are represented over binned BFP expression in (**b**), (**c**), and (**d**), respectively.

**Supplementary Figure 8. Benchmarking Flp systems performance over time using binned mean fluorescence intensity (M.F.I.), fold activation (Fold), and signal-to-noise ratio (SNR).**

(**a**) Legend indicating labels for time-course values. GFP M.F.I., Fold, and SNR are represented over binned BFP expression in (**b**), (**c**), and (**d**), respectively.

**Supplementary Figure 9. Benchmarking ɸC31 systems performance over time using binned mean fluorescence intensity (M.F.I.), fold activation (Fold), and signal-to-noise ratio (SNR).**

(**a**) Legend indicating labels for time-course values. GFP M.F.I., Fold, and SNR are represented over binned BFP expression in (**b**), (**c**), and (**d**), respectively.

**Supplementary Figure 10. Single-cell fluorescence represent tation of selected Cre split systems.**

Single-cell data represented as smoothed scatterplots are plotted of the experiment in **Figure 2c** and **2d**. Axes represent the expression of transfection marker, BFP, in molecules of equivalent fluorescein (MEFL) and GFP output, in MEFL, from recombinase reporters. Density of cytometric events are represented ranging from no events (dark blue) to highest density in dark red. White dots represent single event outliers. Events under a BFP MEFL cutoff of 1x10^6^ (representing non-transfected or poorly transfected cells) were removed from analysis and a red line at GFP MEFL = 1x10^6^ indicates a boundary between cells that are “OFF” (below 1x10^6^) and cells that are “ON” (above 1x10^6^).

**Supplementary Figure 11. Single-cell fluorescence representation of selected Flp split systems.**

Single-cell data represented as smoothed scatterplots are plotted of the experiment in **Figure 2e** and **2f**. Axes represent the expression of transfection marker, BFP, in molecules of equivalent fluorescein (MEFL) and GFP output, in MEFL, from recombinase reporters. Density of cytometric events are represented ranging from no events (dark blue) to highest density in dark red. White dots represent single event outliers. Events under a BFP MEFL cutoff of 1x10^6^ (representing non-transfected or poorly transfected cells) were removed from analysis and a red line at GFP MEFL = 1x10^6^ indicates a boundary between cells that are “OFF” (below 1x10^6^) and cells that are “ON” (above 1x10^6^).

**Supplementary Figure 12. Single-cell fluorescence representation of selected VCre split systems.**

Single-cell data represented as smoothed scatterplots are plotted of the experiment in **Figure 2g** and **2h**. Axes represent the expression of transfection marker, BFP, in molecules of equivalent fluorescein (MEFL) and GFP output, in MEFL, from recombinase reporters. Density of cytometric events are represented ranging from no events (dark blue) to highest density in dark red. White dots represent single event outliers. Events under a BFP MEFL cutoff of 1x10^6^ (representing non-transfected or poorly transfected cells) were removed from analysis and a red line at GFP MEFL = 1x10^6^ indicates a boundary between cells that are “OFF” (below 1x10^6^) and cells that are “ON” (above 1x10^6^).

**Supplementary Figure 13. Single-cell fluorescence representation of selected ɸC31 split systems.**

Single-cell data represented as smoothed scatterplots are plotted of the experiment in **Figure 2i** and **2j**. Axes represent the expression of transfection marker, BFP, in molecules of equivalent fluorescein (MEFL) and GFP output, in MEFL, from recombinase reporters. Density of cytometric events are represented ranging from no events (dark blue) to highest density in dark red. White dots represent single event outliers. Events under a BFP MEFL cutoff of 1x10^6^ (representing non-transfected or poorly transfected cells) were removed from analysis and a red line at GFP MEFL = 1x10^6^ indicates a boundary between cells that are “OFF” (below 1x10^6^) and cells that are “ON” (above 1x10^6^).

**Supplementary Figure 14. Single-cell fluorescence representation of selected TP901 and Bxb1 split systems.**

Single-cell data represented as smoothed scatterplots are plotted of the experiment in **Figure 2k** through **2n**. Axes represent the expression of transfection marker, BFP, in molecules of equivalent fluorescein (MEFL) and GFP output, in MEFL, from recombinase reporters. Density of cytometric events are represented ranging from no events (dark blue) to highest density in dark red. White dots represent single event outliers. Events under a BFP MEFL cutoff of 1x10^6^ (representing non-transfected or poorly transfected cells) were removed from analysis and a red line at GFP MEFL = 1x10^6^ indicates a boundary between cells that are “OFF” (below 1x10^6^) and cells that are “ON” (above 1x10^6^).

**Supplementary Figure 15.** **Comparison of domain orientations with four splits of Flp recombinase.**

(**a**) Flp was split into two fragments at amino acid S and fused to gibberellin-inducible (GAI/GID1), rapalog-inducible (FKBP/FRB) or abscisic acid-inducible (PYL/ABI) dimerization domains. Split recombinases were transfected along with reporters that yield green fluorescence protein (GFP) expression upon site-specific recombination (**b**). Constitutive Flp (black square) or a blank vector (gray square) were transfected to indicate highest or lowest expected GFP expression. Split recombinases were also transfected and drug (colored squares) or no drug (white squares) was added to the cell culture medium after two hours. Depictions of domains indicated the orientation of the domains with respect to the schematic in (**a**). M.F.I. indicates mean fluorescence intensity measured with arbitrary units (a.u.). Errors bars represent arithmetic standard error of the mean of three transfected cell cultures.

**Supplementary Figure 16. An alternative pick of two splits does in Flp recombinase does not yield a functional 2-input protein-based AND gate.**

(**a**) A two-input protein-based AND-gate is created by splitting Flp recombinase at two locations (S_1_ = 168, S_2_ = 396) and fused to the gibberellin (GIB) and abscisic acid (ABA)-associated chemical inducible dimerization domains. (**b**) Plotted results indicate GFP mean fluorescence intensity (M.F.I.) of four conditions of GIB and ABA. Error bars of M.F.I. indicate the arithmetic standard error of the mean between three transfected cell cultures.

**SUPPLEMENTARY FIGURE 1**

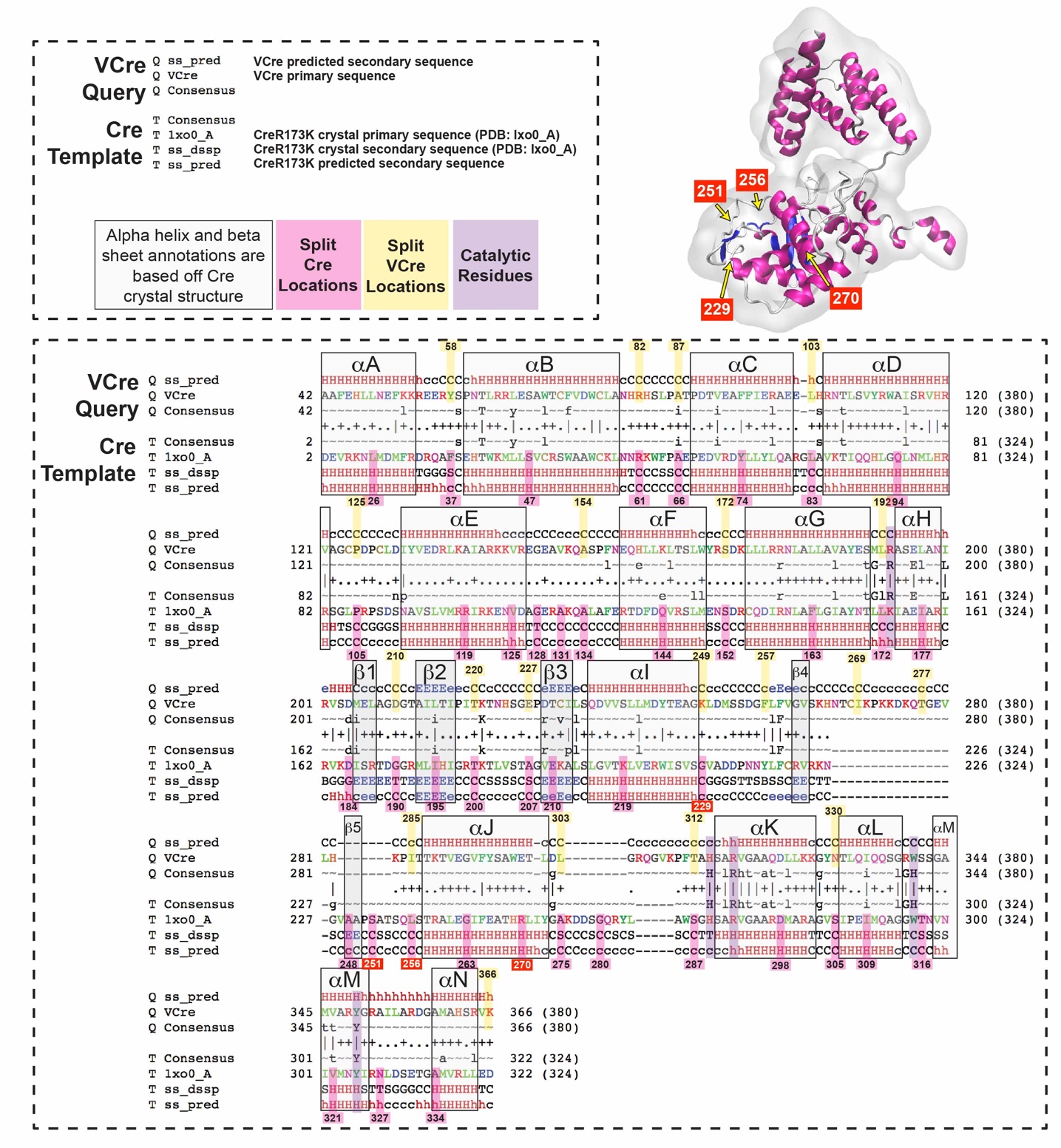

**SUPPLEMENTARY FIGURE 2**

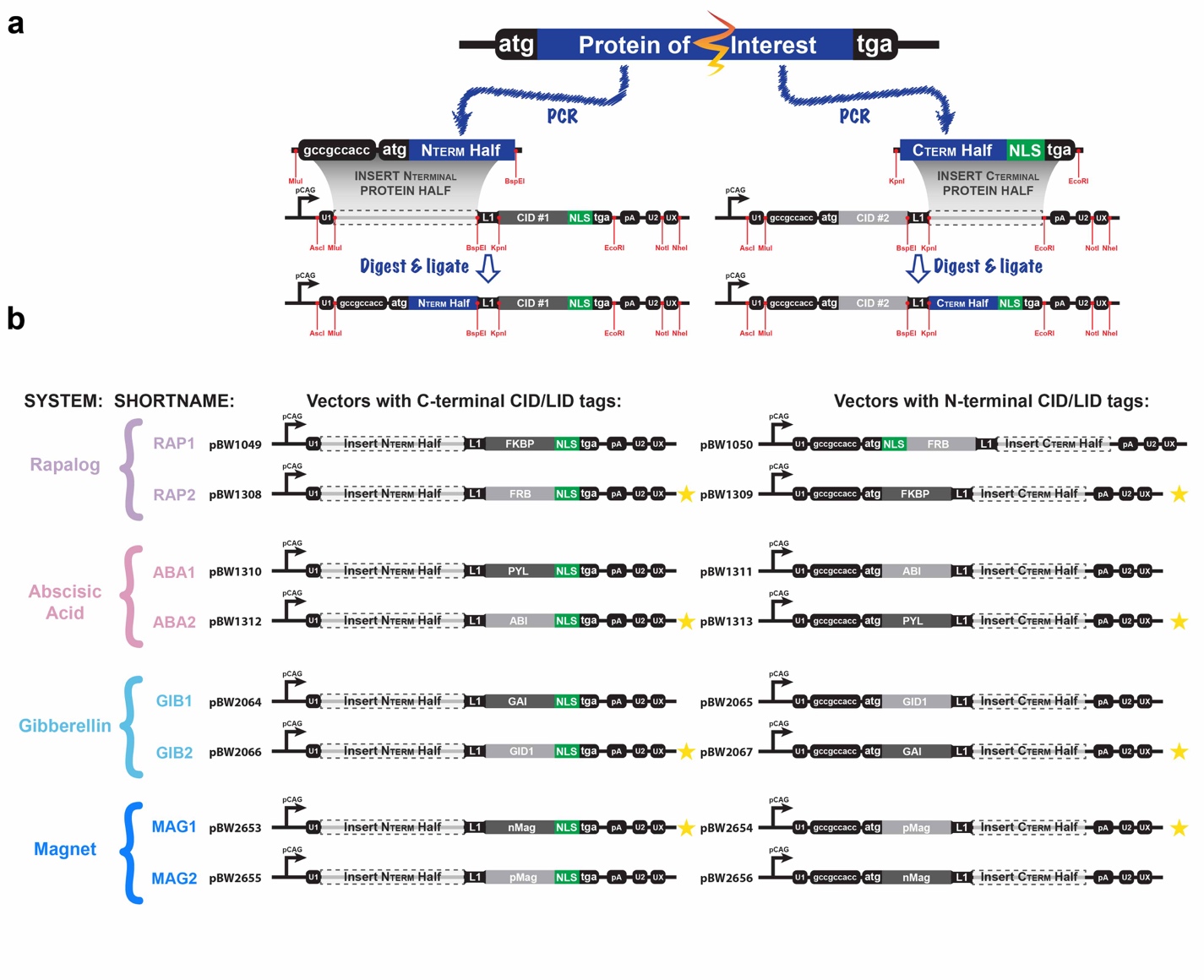
**SUPPLEMENTARY FIGURE 3**

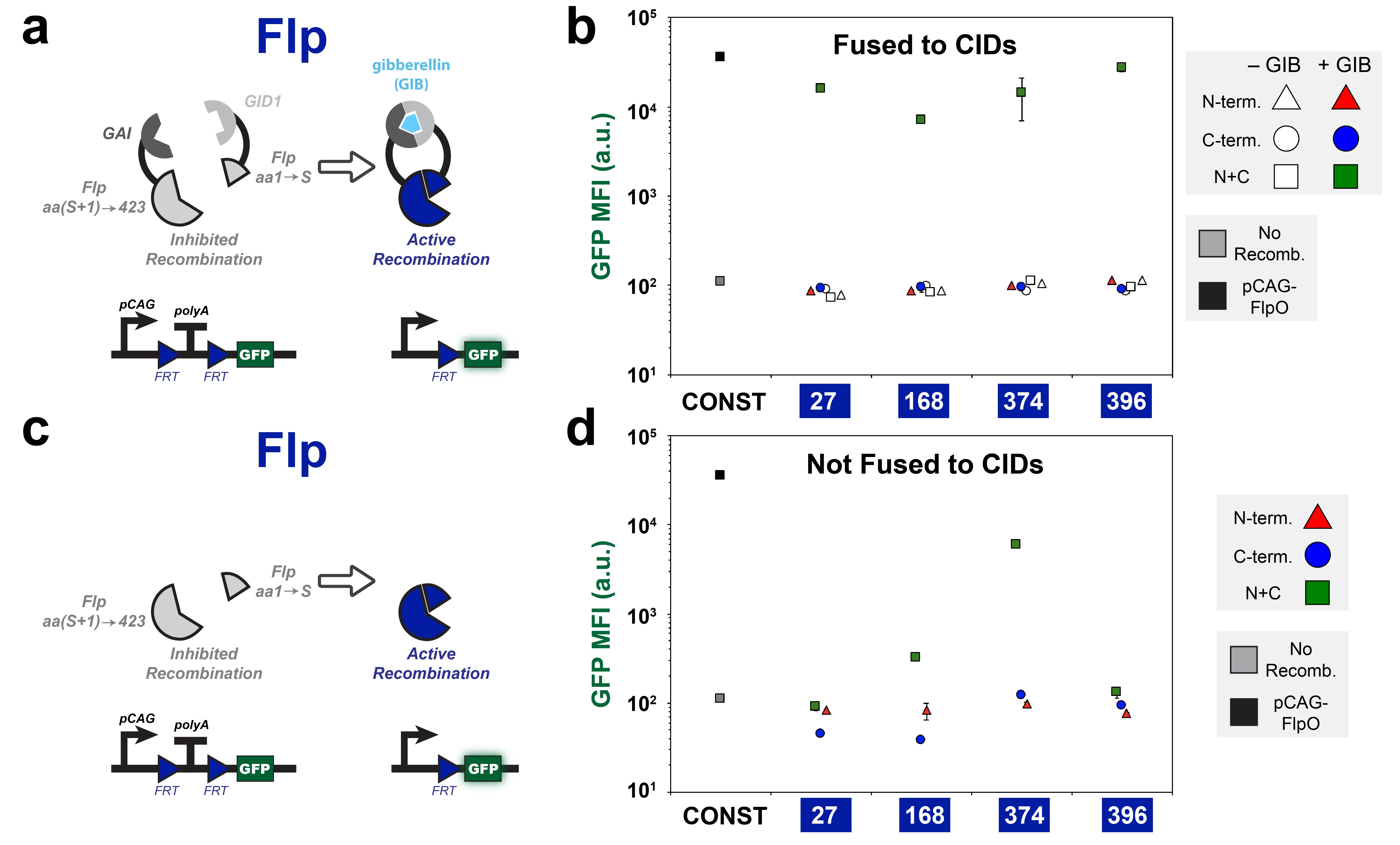
**SUPPLEMENTARY FIGURE 4**

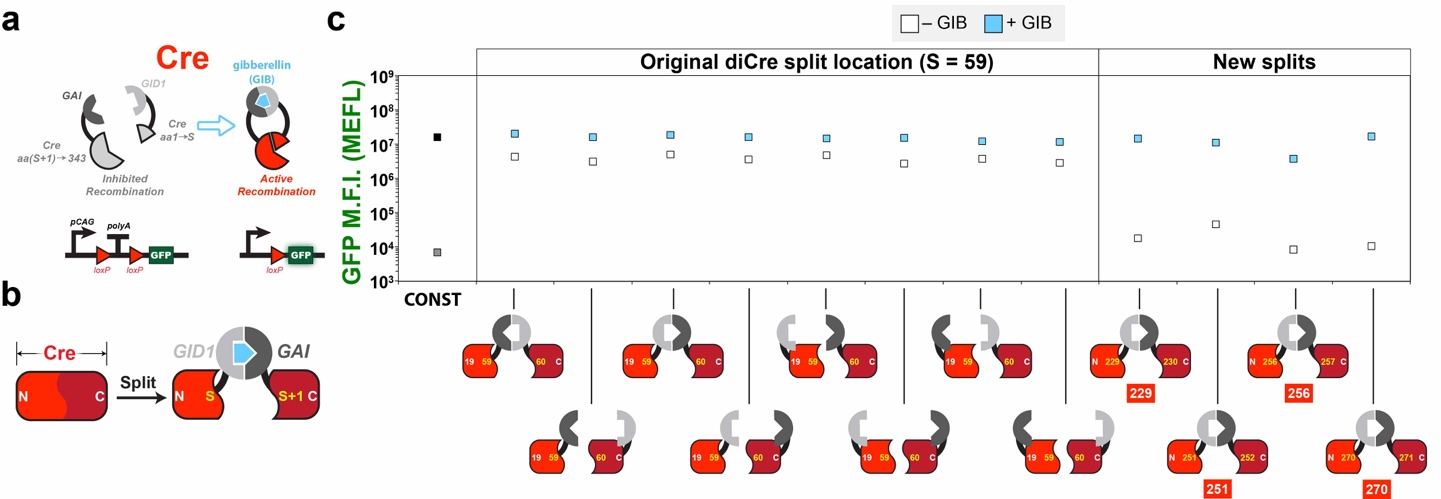

**SUPPLEMENTARY FIGURE 5**

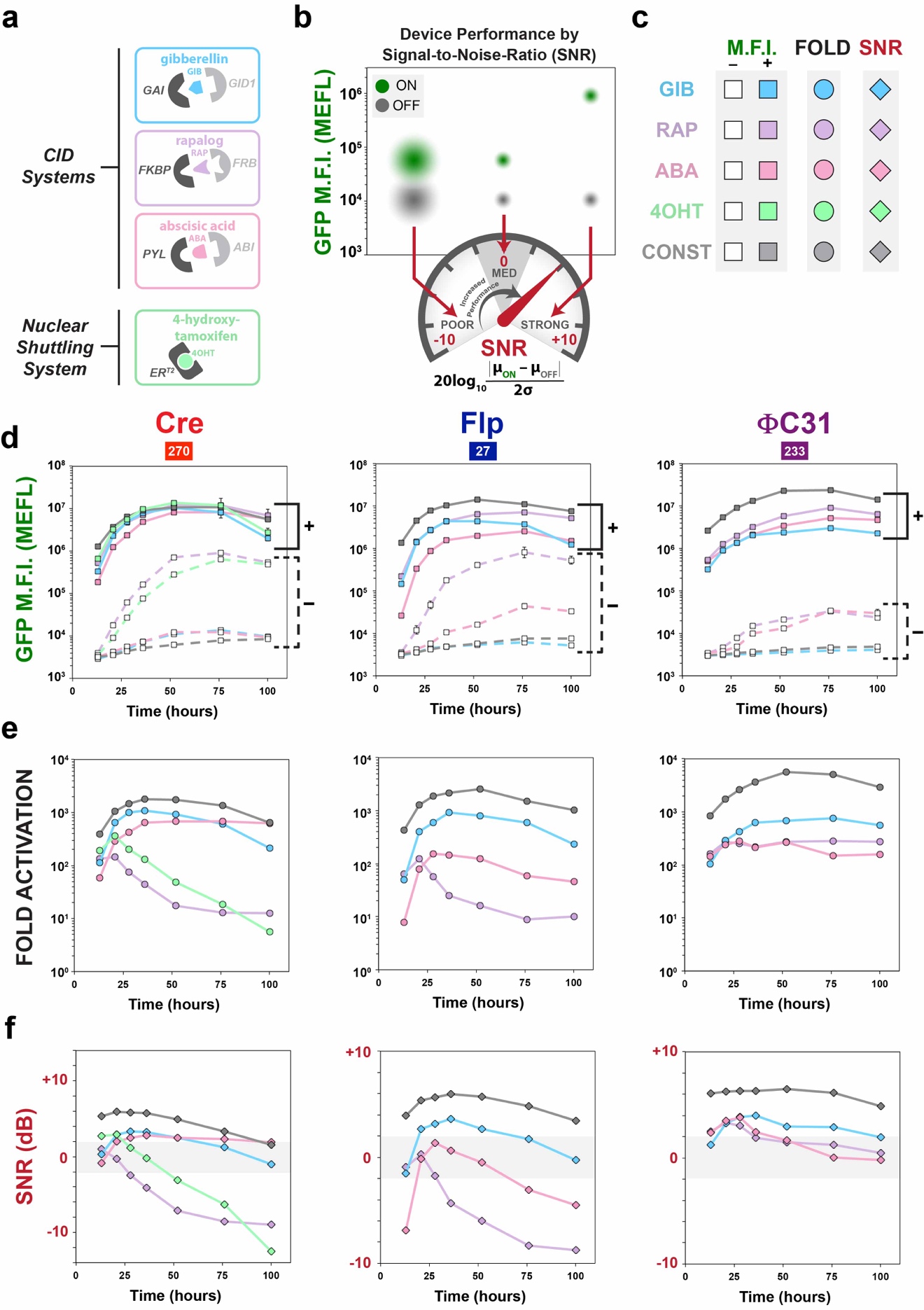
**SUPPLEMENTARY FIGURE 6**

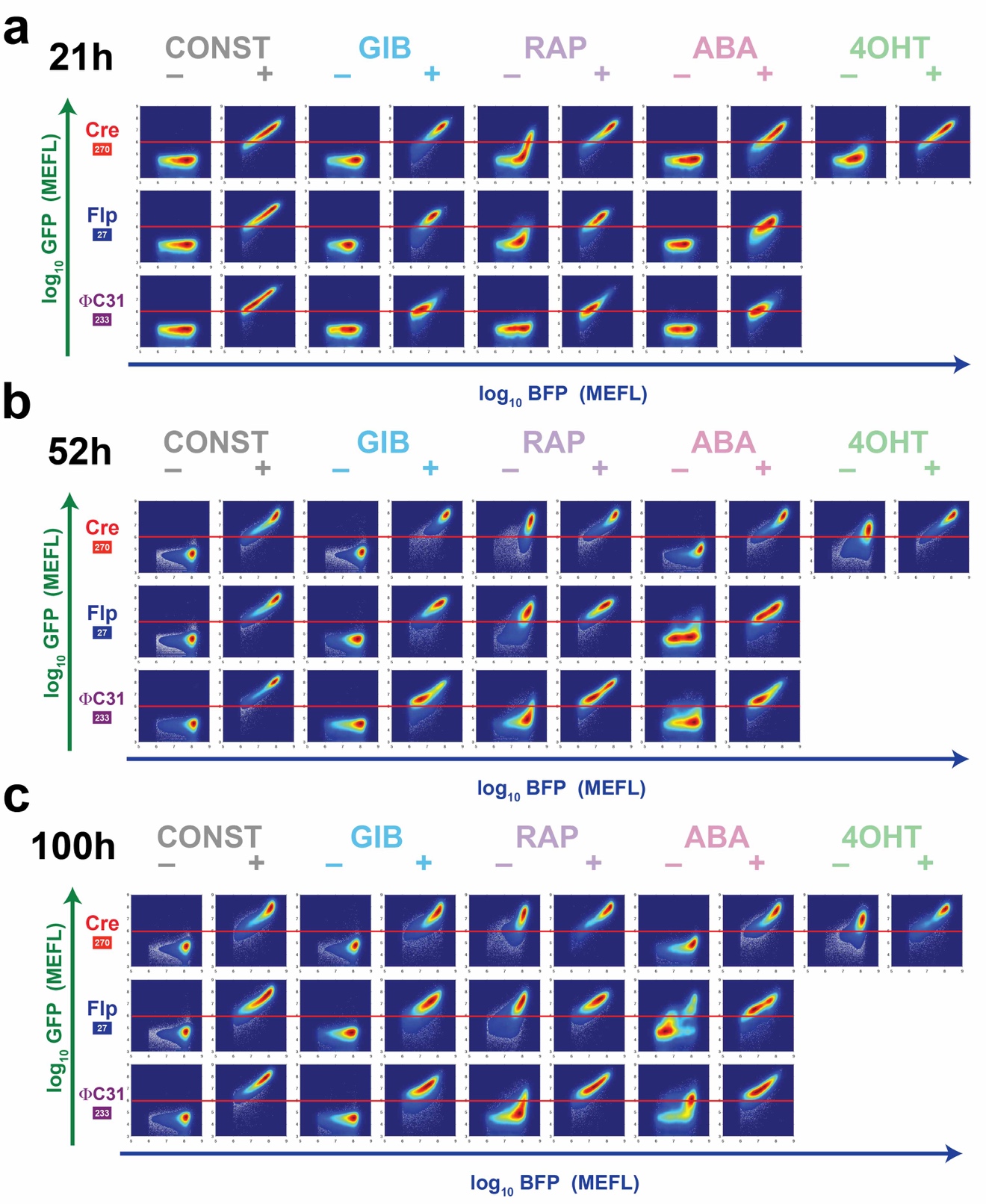
**SUPPLEMENTARY FIGURE 7**

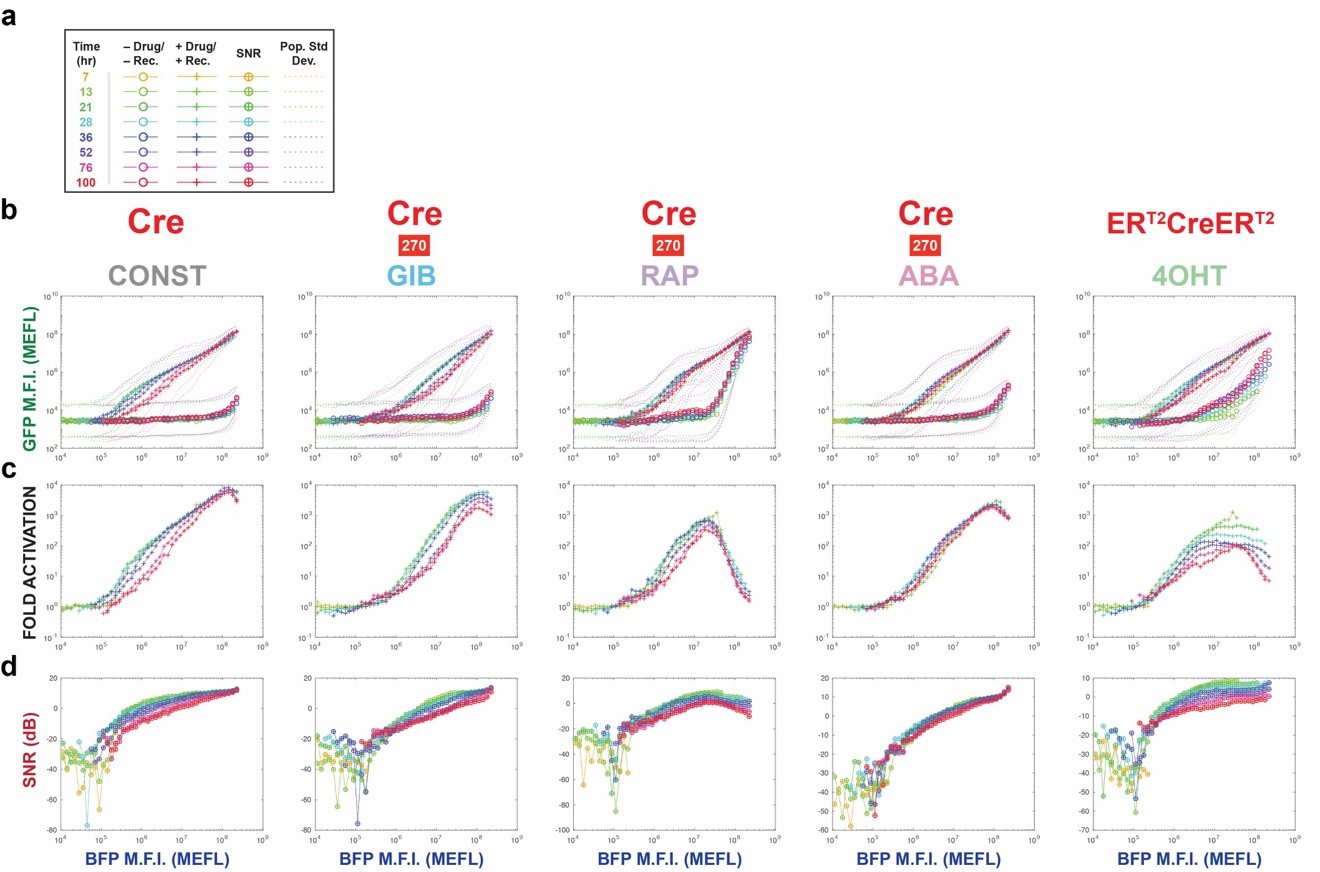

**SUPPLEMENTARY FIGURE 8**

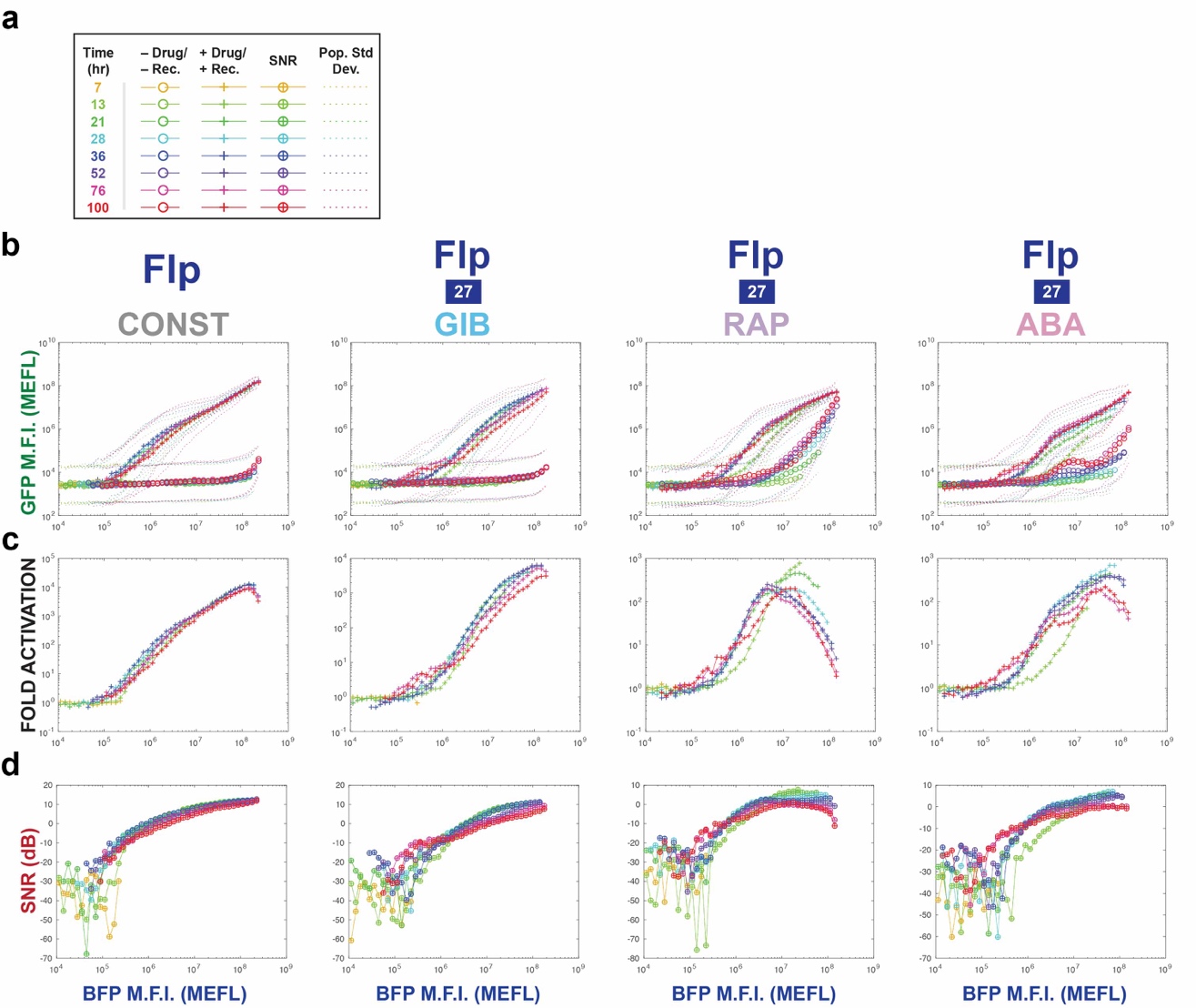
**SUPPLEMENTARY FIGURE 9**

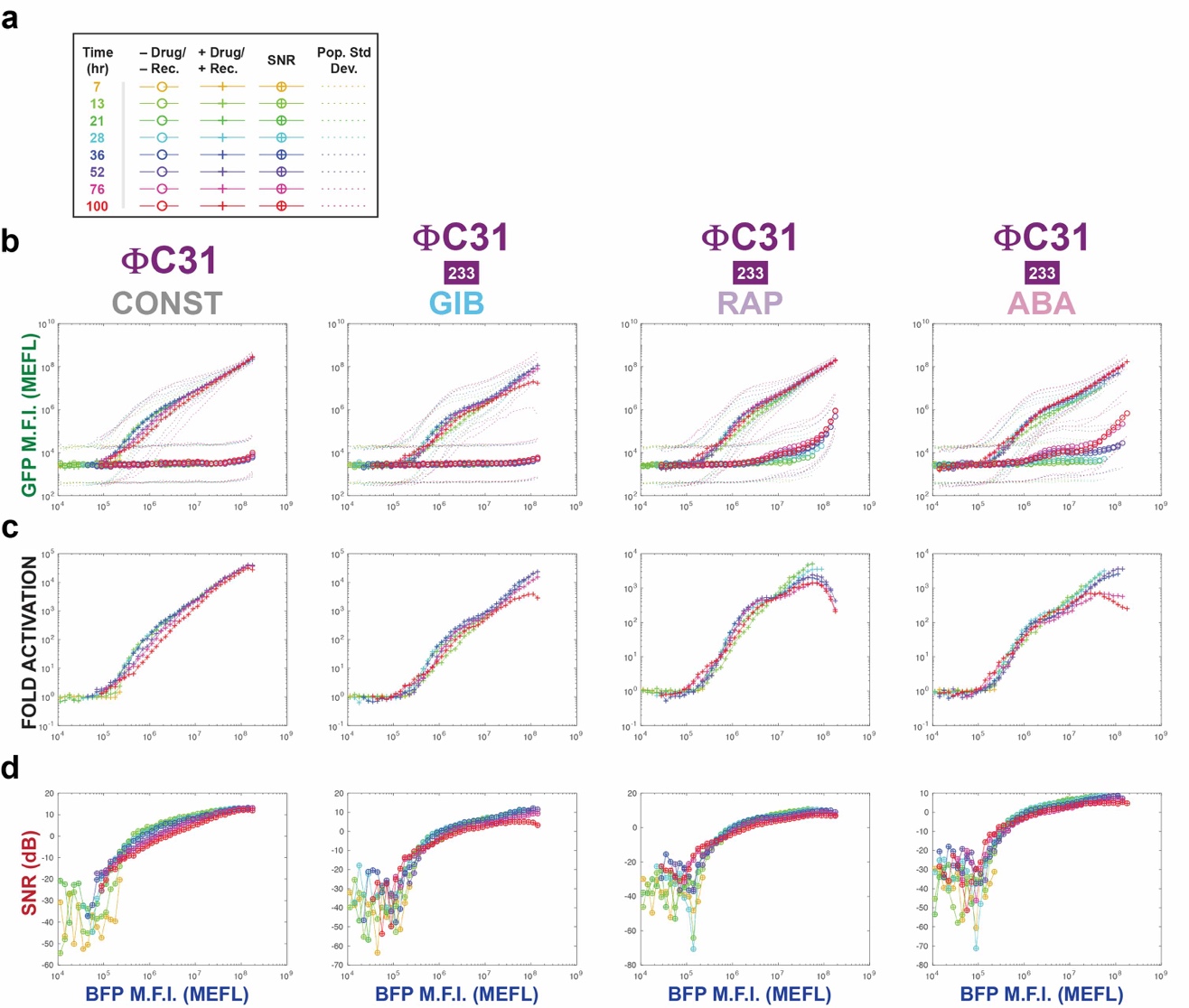
**SUPPLEMENTARY FIGURE 10**

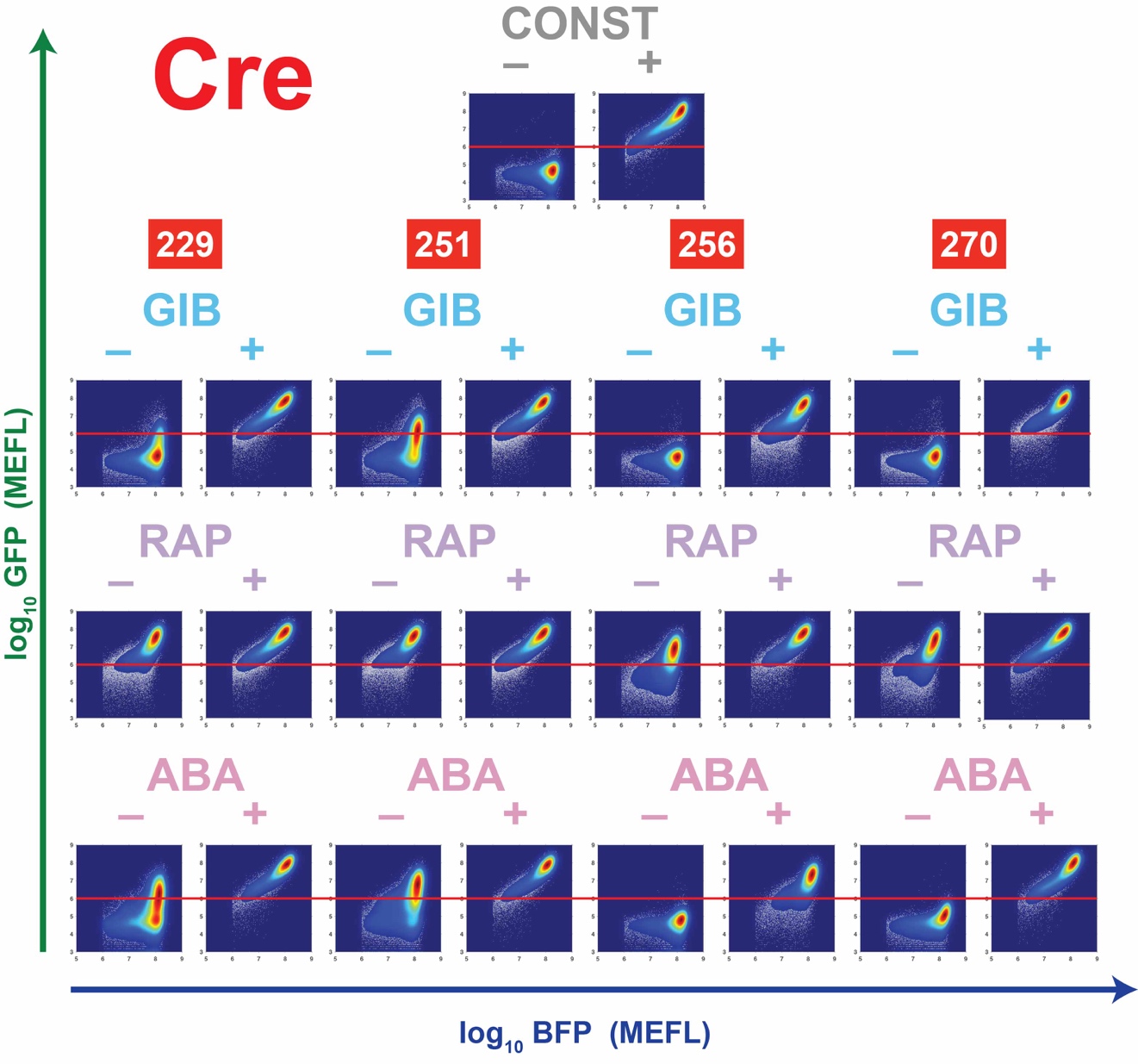

**SUPPLEMENTARY FIGURE 11**

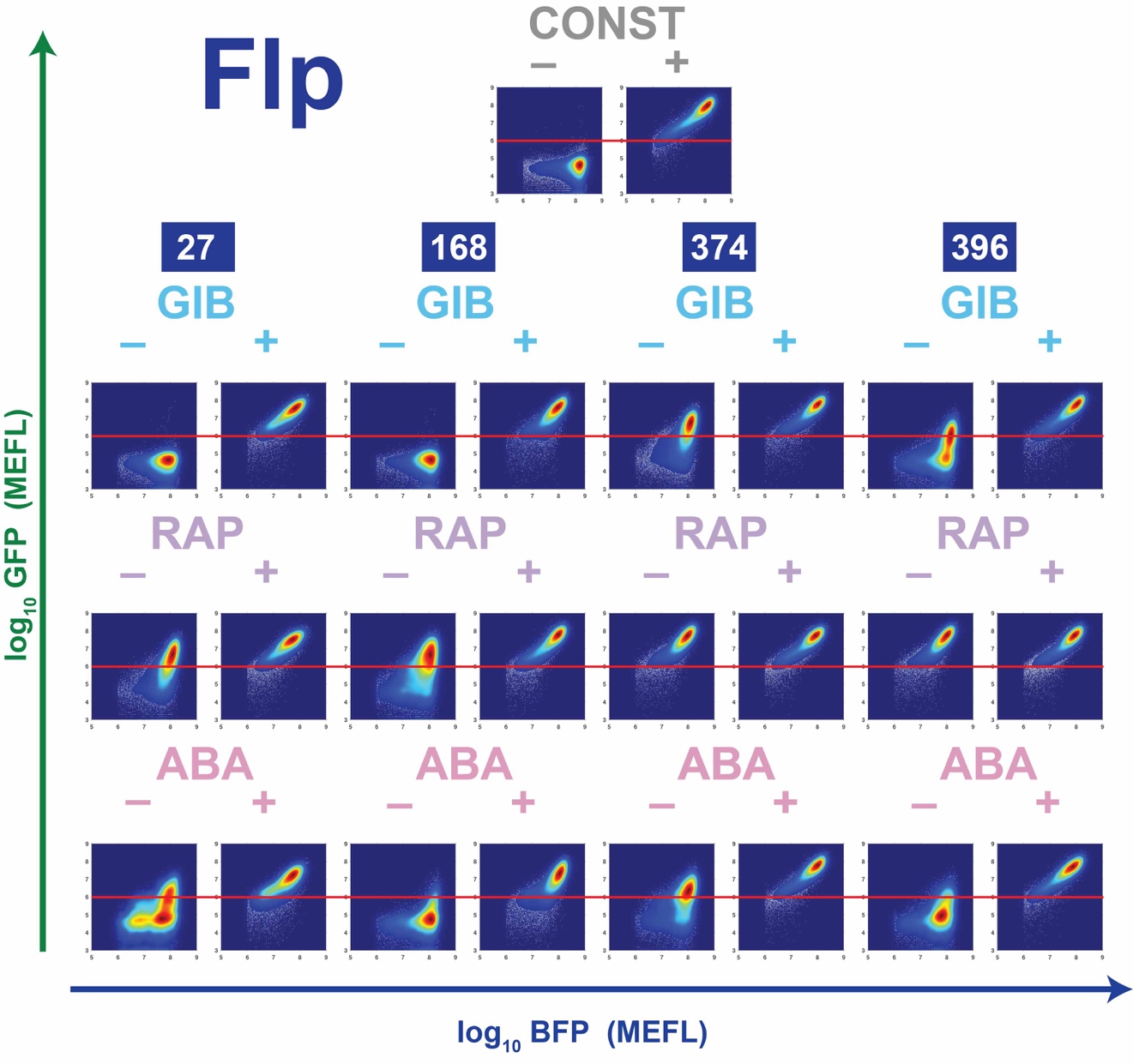

**SUPPLEMENTARY FIGURE 12**

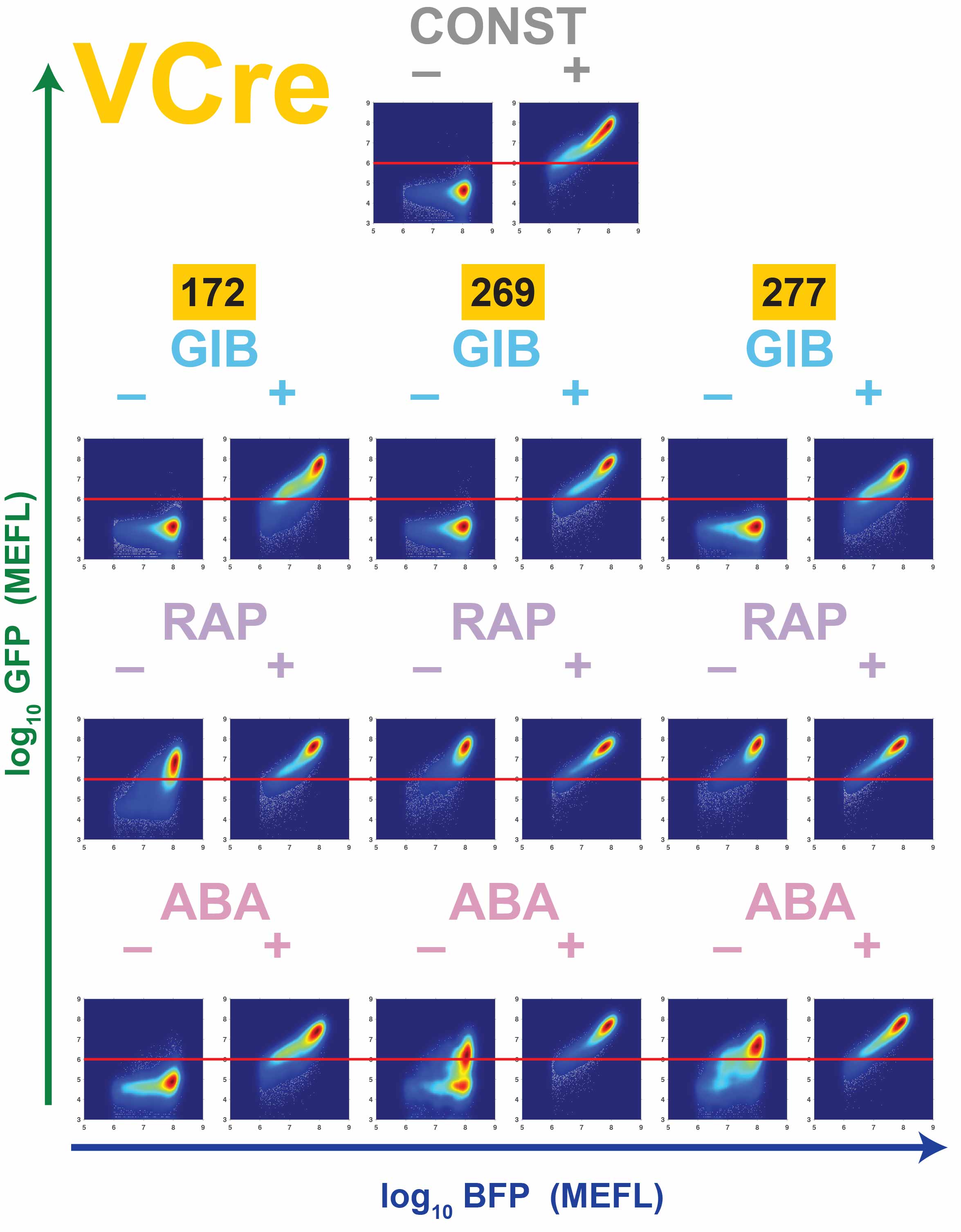
**SUPPLEMENTARY FIGURE 13**

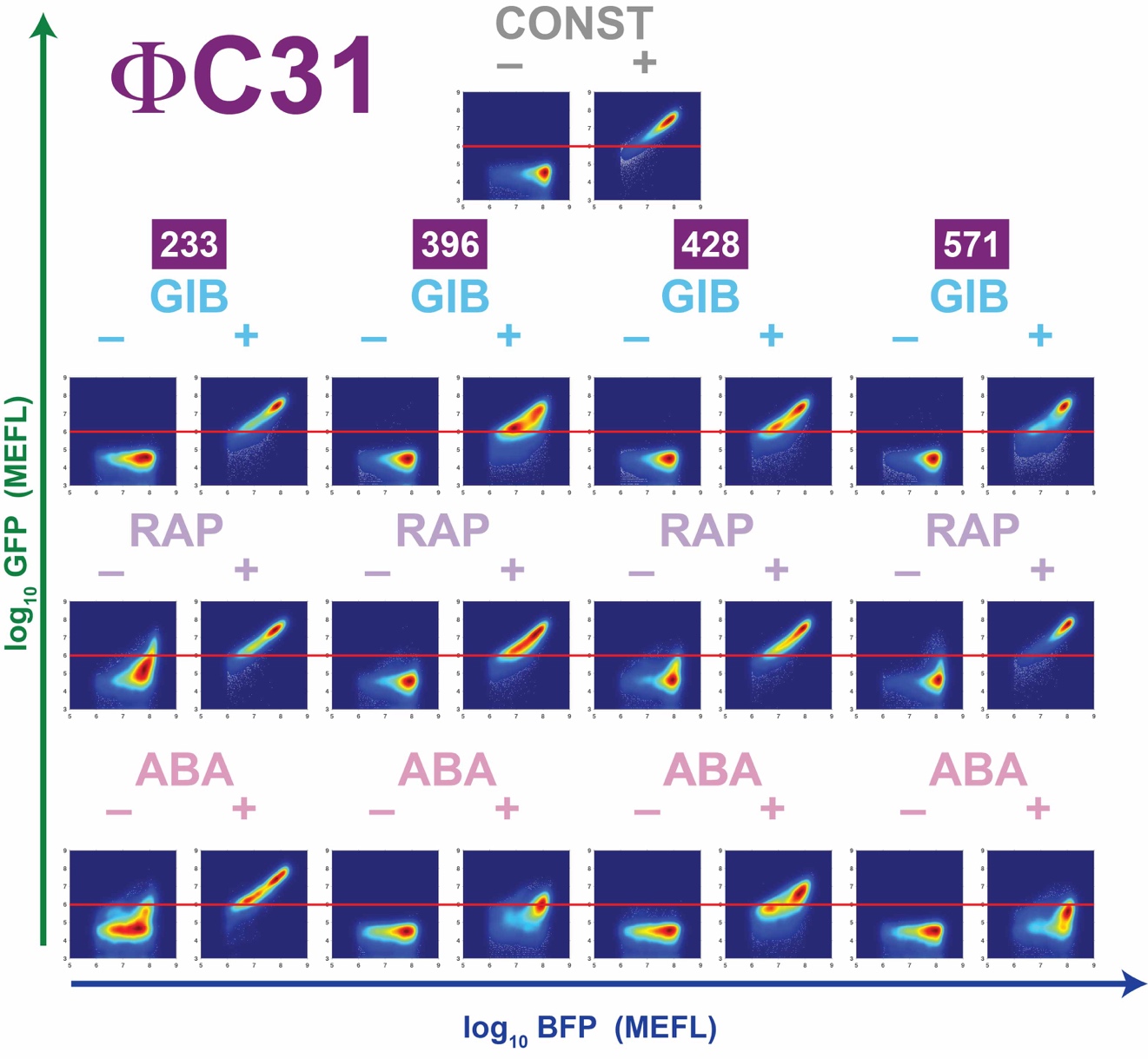
**SUPPLEMENTARY FIGURE 14**

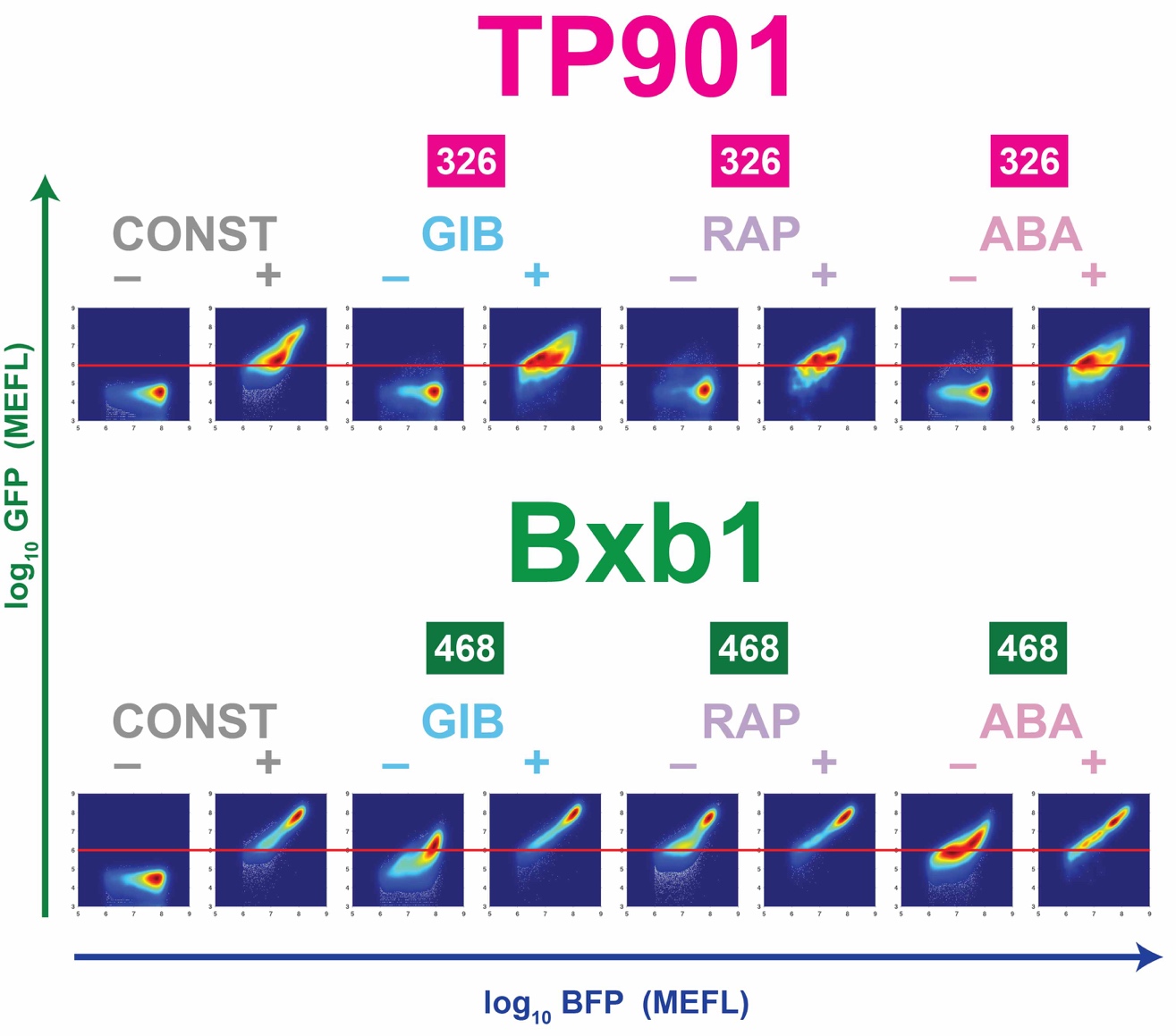
**SUPPLEMENTARY FIGURE 15**

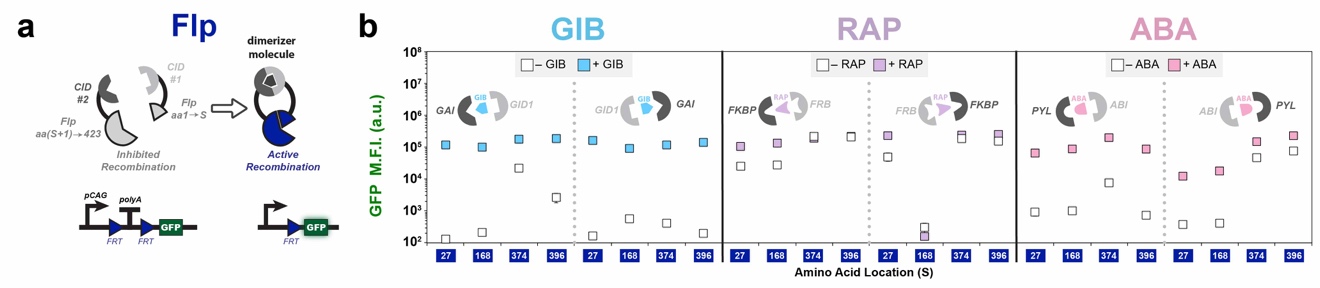

**SUPPLEMENTARY FIGURE 16**

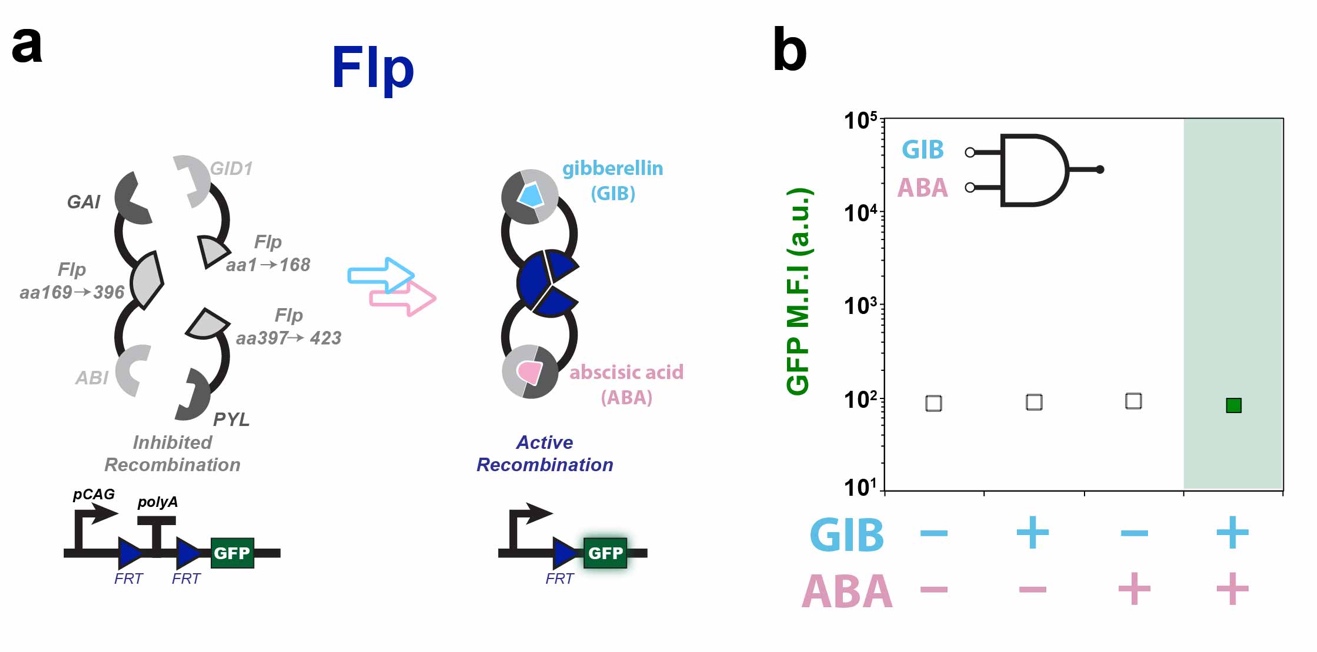
